# Microbial competition reduces interaction distances to the low µm-range

**DOI:** 10.1101/2020.01.22.915835

**Authors:** Rinke J. van Tatenhove-Pel, Tomaž Rijavec, Aleš Lapanje, Iris van Swam, Emile Zwering, Jhonatan A. Hernandez-Valdes, Oscar P. Kuipers, Cristian Picioreanu, Bas Teusink, Herwig Bachmann

**Author notes:** Address correspondence to Herwig Bachmann,.

## Abstract

Metabolic interactions between cells affect microbial community compositions and hence their function in ecosystems. It is well-known that under competition for the exchanged metabolite, concentration gradients constrain the distances over which interactions can occur. However, interaction distances are typically quantified in two-dimensional systems or without accounting for competition or other metabolite-removal, conditions which may not very often match natural ecosystems. We here analyze the impact of cell-to-cell distance on unidirectional cross-feeding in a three-dimensional system with competition for the exchanged metabolite. Effective interaction distances were computed with a reaction-diffusion model and experimentally verified by growing a synthetic consortium of 1 µm-sized metabolite producer, receiver and competitor cells in different spatial structures. We show that receivers cannot interact with producers ∼15 µm away from them, as product concentration gradients flatten close to producer cells. We developed an aggregation protocol and created variants of the receiver cells’ import system, to show that within producer-receiver aggregates even low affinity receiver cells could interact with producers. These results show that competition or other metabolite-removal of a public good in a three-dimensional system reduces the interaction distance to the low micrometer-range, highlighting the importance of concentration gradients as physical constraint for cellular interactions.

## Introduction

Microbial interactions are observed in dense biofilms (0 µm between cells) as well as in oceans (>100 µm between cells), demonstrating that cells interact at various distances [1–4]. These interactions influence the selection pressure within an environment, and therefore affect the structure and evolution of microbial communities [5]. As these communities play an important role in many ecosystems, from global biogeochemical fluxes [6] to human health [7], understanding and controlling these interactions is of high importance.

Metabolites or signaling molecules involved in interactions can be exchanged via contact-dependent and contact-independent transfer mechanisms. Contact-dependent mechanisms require short cell-to-cell distances and use for instance direct contact between cells, vesicle chains or nanotubes for exchange [5]. Contact-independent mechanisms require passive or active transport of the produced compound to the extracellular space, where it subsequently moves via diffusion and convection [5]. Contact-independent interactions can be local (mainly between neighboring cells) or global (within the whole population), depending on the profile of the concentration gradient. *Saccharomyces cerevisiae* for instance uses its extracellular enzyme invertase to split sucrose, resulting in a glucose and fructose gradient around the cell. At high sucrose concentrations both aggregated and single yeast cells can grow (global interactions), while at low sucrose concentrations only aggregated yeast cells grow (local interactions) [8]. A similar pattern is observed for the extracellular protease of *Lactococcus lactis*, which activity results in a peptide gradient around the cell. At high cell densities both protease positive and protease negative cells grow, while at low cell densities mainly protease positive cells grow, since only they can benefit from their produced peptides [9].

Whether contact-independent interactions are local or global depends on the distance between cells and the concentration gradient profile, which is affected by the metabolite source, the metabolite-sink and the diffusion and convection rate between them. The metabolite source can for instance be a producer cell [10], a nutrient pool in the environment [11] or an extracellular enzyme [8, 9]. The metabolite-sink can be a metabolite consuming cell [12, 13], a metabolite degrading enzyme [14] or the volume of the system, as dilution reduces the metabolite concentration [8, 9]. Although the exact nature of the source and sink are often only implicitly mentioned in these studies, their importance is well-known. Costly cooperative interactions are for instance more likely to evolve when cells are close to each other, because cooperators compete with wildtype non-cooperators for the excreted metabolite [5, 15, 16]. Selection for interactions is therefore often done by co-culturing cells on agar plates [17–19], and it is also described that interacting cells evolved aggregating phenotypes [20].

These examples show that in the presence of a metabolite-removing sink, concentration gradients constrain the distances over which interactions can occur. It is however not clear at what distances such interactions occur. Previous studies either quantified these distances in two-dimensional systems [11–13] or without a metabolite-removing sink [13, 14], while natural microbial communities reside in three-dimensional environments in which competing metabolite consumers and other types of metabolite-removing sinks are very likely to be present. We therefore combined computational and experimental analyses to provide a more systematic and quantitative perspective on the impact of cell-to-cell distance on metabolic interactions in three dimensions and in the presence of metabolite-removing sinks. The reaction-diffusion model and experimental results show that in these conditions receiver cells fixed at ∼15 µm from glucose producing cells cannot interact with the producer cells, while producer-receiver aggregation facilitates metabolic interactions even when receiver cells have a low affinity for the product caused by genetic variation of their glucose import systems. These results suggest that competition or other metabolite-removal in a three-dimensional system reduces interaction distances to well below 15 µm.

## Materials and methods

### Strains and media

All the strains that were used are listed in Table 1. *Lactococcus lactis* NZ9000 strains PTSman_GFP, PTScel_GFP, and glcU_GFP were obtained by a single-crossover integration of vector pSEUDO::*Pusp45-gfp* [21] into the *pseudo* 10 locus on the chromosome of *L. lactis* NZ9000Δ*ptcC*Δ*glcU*, NZ9000Δ*ptnABCD*Δ*glcU*, and NZ9000Δ*ptnABCD*Δ*ptcC* [22], respectively. Integration was performed as previously described [23]. Transformants were selected on M17-agar plates supplemented with glucose, sucrose and 5 µg/mL erythromycin.

**Table 1.** Bacterial strains and plasmids used in this study

| *L. lactis* strain | Description | Reference |
| --- | --- | --- |
| NZ9000Δ*ptcC*Δ*glcU* | Derivative of NZ9000 containing a 1254 bp deletion in *ptcC* and a 864 bp deletion in *glcU*. | [22] |
| NZ9000Δ*ptnABCD*Δ*glcU* | Derivative of NZ9000 containing a 1736 bp deletion in *ptnABCD* and a 864 bp deletion in *glcU*. | [22] |
| NZ9000Δ*ptnABCD*Δ*ptcC* | Derivative of NZ9000 containing a 1736 bp deletion in *ptnABCD* and a 1254 bp deletion in *ptcC*. | [22] |
| MG5267 | *L. lactis* MG1363 with the lactose operon integrated into the genome. | [26] |
| NZ9000 Glc-Lac+ | NZ9000Δ*glk*Δ*ptnABCD* containing a 657-bp deletion in *ptcBA*, carrying pMG8020 (lactose mini-plasmid of 23.7 kb, containing *lacFEGABCD*, derivative of pLP712). | [27] |
| MG1363_GFP | MG1363 carrying pSEUDO::*P<sub>usp45</sub>-gfp*. | [21] |
| NZ9000_PTSman_GFP | Ery<sup>r</sup>, NZ9000Δ*ptcC*Δ*glcU* carrying pSEUDO::*P<sub>usp45</sub>-gfp*. | This work |
| NZ9000_PTScel_GFP | Ery<sup>r</sup>, NZ9000Δ*ptnABCD*Δ*glcU* carrying pSEUDO::*P<sub>usp45</sub>-gfp*. | This work |
| NZ9000_glcU_GFP | Ery<sup>r</sup>, NZ9000Δ*ptnABCD*Δ*ptcC* carrying pSEUDO::*P<sub>usp45</sub>-gfp*. | This work |
| Plasmids | Description | Reference |
| pSEUDO::*P<sub>usp45</sub>-gfp* | Ery<sup>r</sup>, integration vector, pSEUDO::*P<sub>usp45</sub>-sfGFP(Bs)* derivative, carrying the gene coding for the green fluorescent protein (Dasher-GFP). | [21] |

*L. lactis* was grown in chemically defined medium (CDM) described by Otto *et al.* [24], with the following changes: 0.6 g/L NH_4_-citrate, 2.5 mg/L biotin, 0.02 mg/L riboflavin and no folic acid. *L. lactis* NZ9000 Glc-Lac+ was pre-cultured in CDM + 0.95 wt% lactose, *L. lactis* MG5267 in CDM + 0.5 wt% lactose and *L. lactis* MG1363 [25], *L. lactis* MG1363_GFP, *L. lactis* NZ9000_GFP_PTSman, *L. lactis* NZ9000_GFP_glcU and *L. lactis* NZ9000_GFP_PTScel in CDM + 0.5 wt% glucose. Agarose beads contained CDM + 0.4 wt% carbon source and were incubated surrounded by oil, or by CDM + 0.2 wt% carbon source. Mono-cultures were incubated with the same carbon source as their pre-culture, co-cultures were incubated in presence of lactose. Incubations were done at 30°C.

### Aggregation protocol

Producer and receiver cell pre-cultures (10 mL) were washed three times with 0.9% sodium chloride. Receiver cells were resuspended in 2 mL 0.225% sodium chloride, producer cells were resuspended in 0.9% sodium chloride and diluted to an OD_600_ of 1.1. Both were incubated in an ultrasonification bath (Branson 200 Ultrasonic cleaner, Branson Ultrasonics, Danbury, CT, USA) at 46 kHz for 3 minutes to ensure complete resuspension to single cells. The surface of (non-)producer cells was charged positively by electrostatic deposition of polyethyleneimine (PEI; Mr 600 000 - 1 000 000; ∼50% in H_2_O; Sigma-Aldrich, Saint Louis, MO, USA) as follows. Sonicated producer cells were mixed with 0.25% PEI (hydrated, pH 7) in a 1:1 (v/v) ratio and incubated at room temperature for 5 minutes. After incubation cells were collected by centrifugation (900 g, 3 minutes) and washed by replacing the supernatant with 0.9% sodium chloride five times without resuspending the pellet. Washed cells were resuspended in 200 µL 0.9% sodium chloride and sonicated as described above. The surface of washed receiver cells was negatively charged and therefore not further modified [28]. Cell concentrations in the prepared producer and receiver suspensions were measured with flow cytometry (Accuri C6, BD Biosciences, San Jose, CA, USA).

Aggregates were formed electrostatically by mixing the positively charged producer cells with the negatively charged receiver cells, such that the oppositely charged cells stuck to each other. The suspension with negatively charged receiver cells was mixed using a T10 basic ULTRA TURRAX homogenizer with an S10N-5G dispersing element (IKA, Staufen, Germany) at 8000 rpm for 15-20 minutes. While mixing, the positively charged producer cells were added to the negatively charged receiver cells using a 1 mL syringe (Terumo, Tokyo, Japan), a Chemyx Fusion 200 syringe pump (125-400 µL/h, Chemyx Inc., Stafford, TX, USA) and polyethylene tubing (inner diameter 0.38 mm, BD, Franklin Lakes, NJ, USA).

The mixing time (15-20 minutes) and syringe pump flow rate (125-400 µL/h) were adjusted within the mentioned ranges such that the final aggregate percentage was ∼3%.

### Agarose beads formation and analysis

Agarose beads in oil were made by mixing a water and an oil phase. The oil phase contained Novec HFE 7500 fluorinated oil (3M, Maplewood, MN, USA) and 0.2% PicoSurf 1 surfactant (Sphere Fluidics, Cambridge, UK). The water phase contained CDM, 1 wt% melted agarose with ultra-low gelling temperature (Type IX-A, A2576, Sigma-Aldrich, Saint Louis, MO, USA) and cells, and it was prepared as follows. Pre-cultures were washed with phosphate buffered saline (PBS) and the OD_600_ was measured to determine the cell concentration (assuming OD 1 = 10^9^ cells/mL). The total cell concentration in the aggregate suspension was determined using flow cytometry (Accuri C6). The producer cell or aggregate concentration in CDM with agarose was set to 2.7·10^6^/mL, the receiver cell concentration to 8.9·10^7^cells/mL.

300 µL water phase and 700 µL oil phase were mixed using a T10 basic ULTRA TURRAX homogenizer with an S10N-5G dispersing element at 8000 rpm for 5 minutes. Emulsions were subsequently placed on ice for at least 20 minutes, to solidify the agarose beads. After solidification cells could not move and growth therefore resulted in micro-colony formation within the agarose bead. Formed agarose beads had an average diameter of 37 µm (supplementary information, section 1). Based on this average diameter, each bead contained on average 8 receiver cells. In addition to the receivers, ∼2% of the beads contained two or more producer cells/aggregates and ∼19% contained one producer cell/aggregate (∼79% contained no producer/aggregate).

To incubate agarose beads in CDM, 1 mL CDM and 1 mL perfluorooctanol (PFO, Alfa Aesar, Ward Hill, MA, USA) were added to the emulsion after solidification. This leads to the breaking of the emulsion and separation of the water and oil phase upon gently mixing. Subsequently the water phase, containing agarose beads in CDM, was separated from the oil phase and incubated while rotating. For incubation in presence of competing glucose-consumers 10^9^ *L. lactis* MG1363 cells per mL were added to the CDM surrounding the agarose beads (supplementary information, section 2).

Growth in agarose beads was analyzed with flow cytometry (Accuri C6). Agarose beads in CDM were measured directly. For agarose beads in oil the emulsion was first broken by adding 240 µL PBS and 300 µL PFO to 60 µL emulsion, followed by gently mixing. The water phase, containing agarose beads in PBS, was separated from the oil phase and measured using flow cytometry. Details about the flow cytometry gating strategy and data analysis are shown in supplementary information section 3.

### Three-dimensional reaction-diffusion model

A three-dimensional, numerical reaction-diffusion model was implemented in COMSOL Multiphysics (COMSOL 5.0, Comsol Inc., Burlington, MA, USA). Two spherical agarose beads were placed in a cubic computational domain. One bead contained a producer cell that secreted glucose with a constant rate, and both beads contained eight receiver cells that consumed glucose based on Monod (saturation) kinetics. The concentration at the agarose bead surface resulted from a partition coefficient which was set to 0 to model incubation in oil, and to 1 to model incubation in CDM. The diffusion coefficient of glucose was set to 6.7·10^-10^ m^2^/s [29] both inside and outside agarose beads [30], and 10 times lower in micro-colonies [29, 31]. A time-dependent study yielded the spatial distribution of glucose. See supplementary information section 4 for more details about the geometry and used parameters.

## Results

### Reaction-diffusion modelling predicts short interaction distances in three-dimensional systems

To compare concentration gradients in two- and three-dimensional systems we made reaction-diffusion models in COMSOL Multiphysics (supplementary information, section 4.2). The concentration gradients around a producer cell were calculated either in cube to mimic a three-dimensional system, or in a thin plate to mimic a two-dimensional system (plate thickness of 1.1 µm, roughly matching the producer cell diameter of 1 µm). In both cases the total volume was 1 nL (10^6^ cells/mL). The model predicted that in the thin plate the maximal concentration is halved at 24 µm from the producer cell, while in the cube this distance is 0.6 µm (Figure S5). This indicated that in three-dimensional systems the distances at which cells can interact are significantly shorter than in two-dimensional systems.

### Design of a synthetic consortium and three-dimensional spatial structure for growth

To study how concentration gradients constrain interactions between micro-organisms in a three-dimensional environment, we extended the cubic model to contain producer and receiver cells, and analyzed the impact of cell-to-cell distance on the interaction (supplementary information, section 4.3). To experimentally validate the model results we constructed synthetic consortia using four *L. lactis* strains. 1) A “producer” that takes up lactose and hydrolyzes it intracellularly to glucose and galactose. It was engineered to not metabolize glucose, which was therefore secreted while the cells grew on galactose. 2) A GFP-expressing “receiver” that can take up and grow on glucose, but not lactose. 3) A “non-producer” that takes up lactose. It uses both the glucose and galactose moiety for growth, and therefore does not secrete glucose. 4) A “competing glucose-consumer” (Figure 1A). To co-culture these cells in a three-dimensional system, glucose-producers and -receivers (the unidirectional cross-feeders) were encapsulated in solidified agarose beads with an average diameter of ∼40 µm. For negative controls, glucose-producers were replaced by glucose-“non-producers”. Cells were embedded in the beads either as separate cells (∼15 µm between cells) or as aggregates (0 µm between cells) (Figure 1B). During incubation agarose beads were separated either by oil or by CDM (Figure 1C). Separation by oil prevented diffusion of glucose from beads, enabling us to validate that cells can grow and interact in agarose beads. Separation by CDM resulted in glucose diffusion from beads, enabling us to study unidirectional cross-feeding in presence of a concentration gradient in a three-dimensional system. To investigate the effect of metabolite-removal on the interaction distances, interactions were analyzed in presence and absence of competing glucose-consumers outside the beads (Figure 1C).

**Figure 1.**
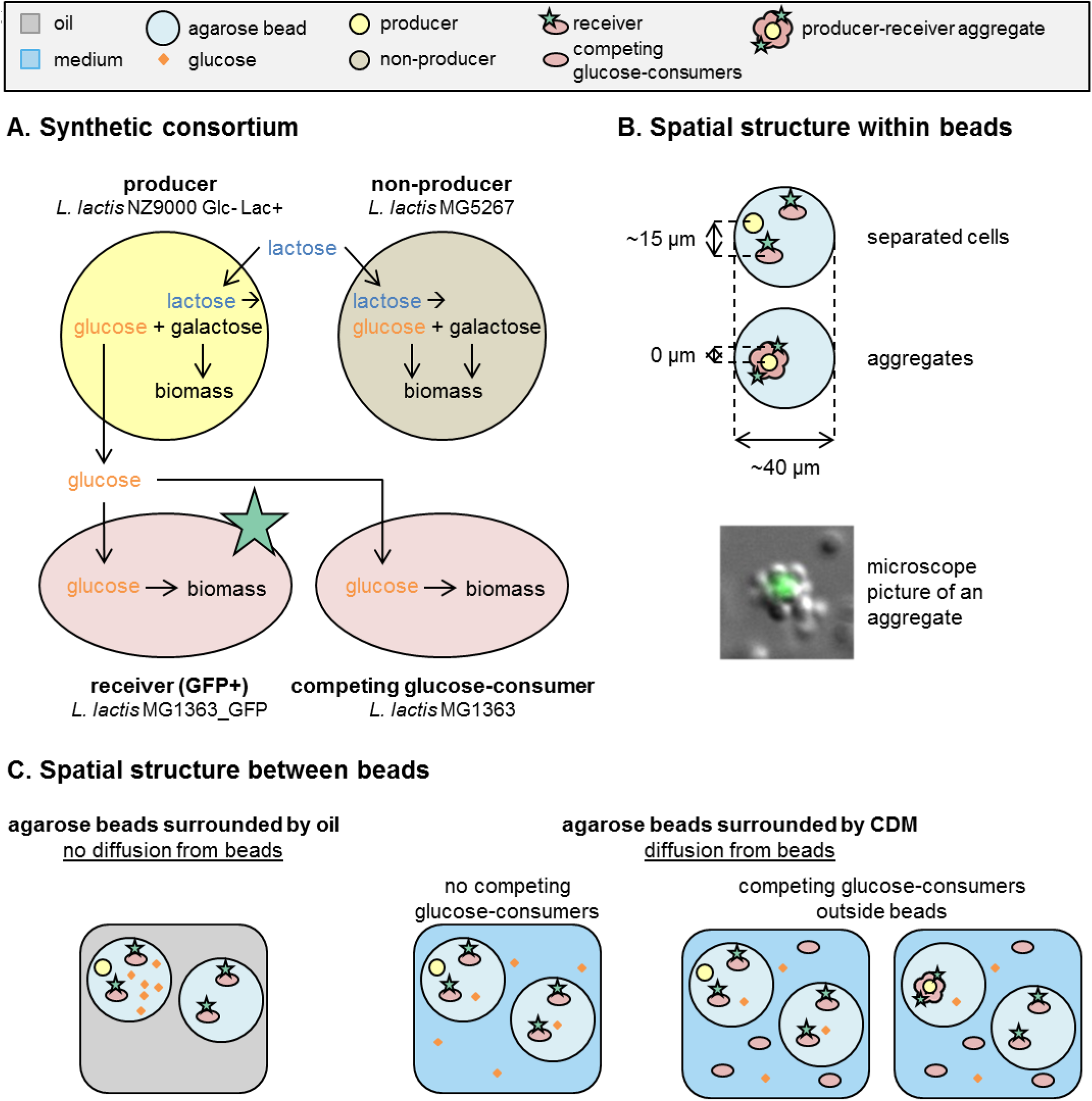
Metabolic interactions in three-dimensional spatially structured environments. **(A)** The four *L. lactis* strains that were used to make synthetic consortia: 1) “producers” which take up lactose and secrete glucose, 2) “receivers” which take up glucose and express GFP, 3) “non-producers” which take up lactose but do not secrete glucose and 4) “competing glucose-consumers” which take up glucose (supplementary information, section 2). **(B)** The three-dimensional spatial structure within agarose beads. A distance of 15 µm between cells is comparable to a homogeneous distribution of 3·10^8^ bacteria/mL. For visibility reasons the microscope picture shows a GFP-expressing cell surrounded by non-fluorescent cells. Aggregates used in experiments were formed oppositely: a (non-)producer cell was surrounded by GFP-expressing receivers. **(C)** The three-dimensional spatial structure between agarose beads. Aggregates were only incubated in presence of competing glucose-consumers.

To analyze if we could detect growth in agarose beads we cultured producers and receivers in beads surrounded by oil (no glucose diffusion from beads) and analyzed the beads with flow cytometry before and after incubation. Beads inoculated with mono-cultures of producers (lactose as carbon source) or receivers (glucose as carbon source) showed an increased forward scatter after incubation, indicating that cells could grow inside beads. In agreement with our expectation the fluorescence/scatter ratio of beads with growth was low for beads with producers and high for beads with GFP-expressing receivers (Figure 2B).

**Figure 2.**
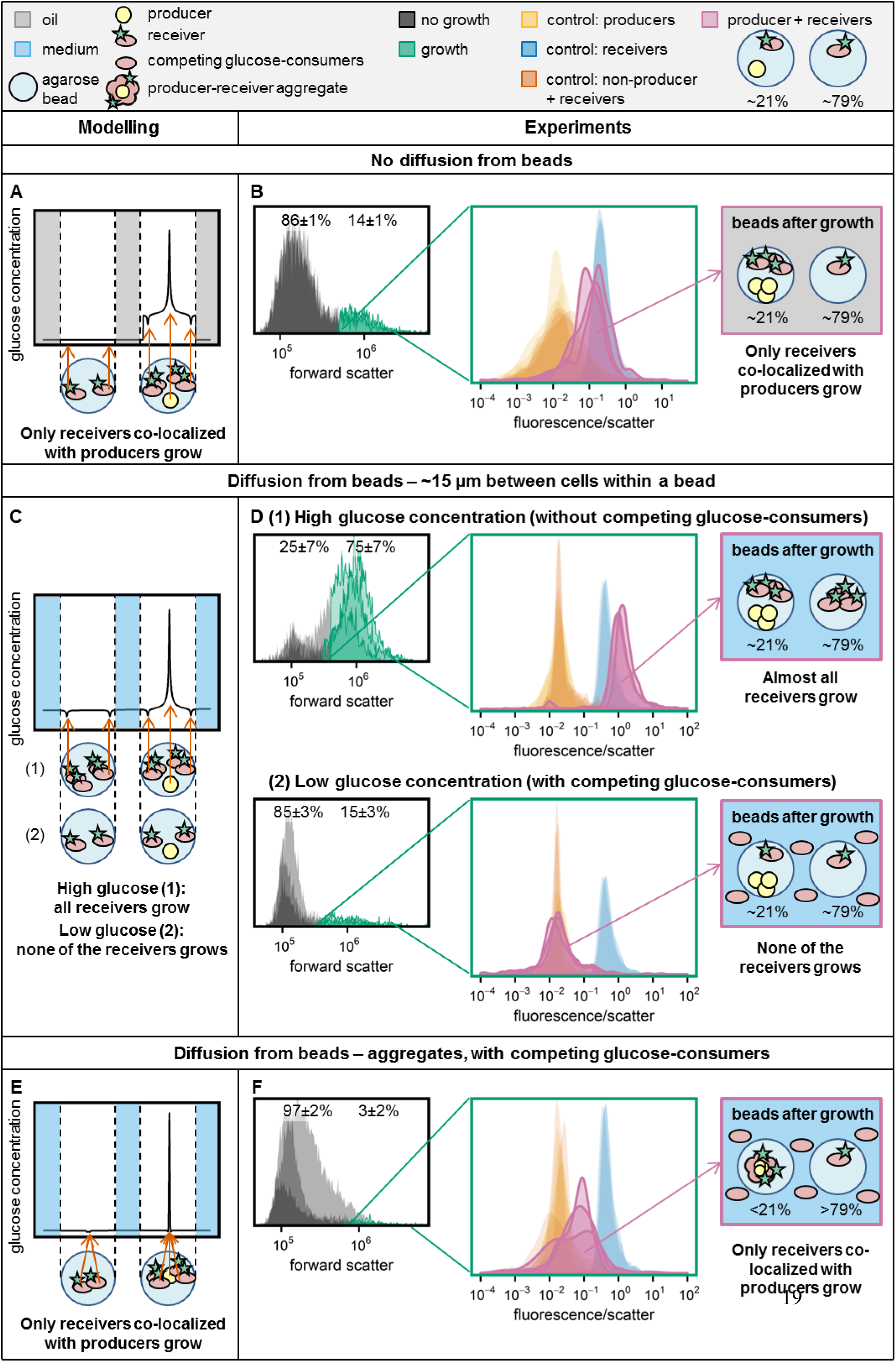
Consortium response in different spatial structures. Panels **(A)**, **(C)**, and **(E)** show the predicted concentration gradient at the diagonal of the cube, for the following spatial structure: (A) No diffusion from beads, (C) Diffusion from beads, ∼15 µm between cells within a bead and (E) diffusion from beads, aggregated cells within a bead. In panels **(B)**, **(D)** and **(F)** the experimental results are shown for these different spatial structures. The forward scatter histograms show the populations that were gated as “growth” in the producer-receiver co-cultures (n=3). From these populations the fluorescence/scatter signal was determined. Next to the producer-receiver co-culture control samples were included: receivers only, producers only and co-cultures of non-producers and receivers (n=3 for each of them). The non-producers and receivers, and the producers only controls are overlapping in all plots. The schematic drawing at the right is based on the experimental data and shows which cells could grow in producer-receiver co-cultures.

To validate that cells could also interact within beads, we made beads such that ∼21% of them contained both a producer and receivers and ∼79% contained receivers only. Because the metabolic interaction is unidirectional, we expected that in presence of lactose the producers would always grow, while the receivers would only grow when glucose, secreted by producers, was available to them (Figure 2A). After incubation 14±1% of the agarose beads showed an increased forward scatter, and these beads had a high fluorescence/scatter ratio (Figure 2B, Table 2). This is close to the expected 21% of beads with producer and receivers, indicating that receivers could only grow in beads with producers.

**Table 2.** Consortium response in different spatial structures. Summary of the experimental results of Figure 2.

| incubation condition | cell-to-cell distance | producer + receivers (~21%)<br>receivers only (~79%) |  | non-producer + receivers (~21%)<br>receivers only (~79%) |  |
| --- | --- | --- | --- | --- | --- |
|  |  | beads with growth | fluorescence/<br>scatter ratio | beads with growth | fluorescence/<br>scatter ratio |
| no diffusion from beads | ~15 $\mu\text{m}$ | $14 \pm 1\%$ | High | $16 \pm 0\%$ | Low |
| diffusion from beads,<br>no competing glucose<br>consumers | ~15 $\mu\text{m}$ | $75 \pm 7\%$ | High | $6 \pm 1\%$ | Low |
| diffusion from beads,<br>competing glucose<br>consumers | ~15 $\mu\text{m}$ | $15 \pm 3\%$ | Low | $9 \pm 2\%$ | Low |
| | 0 $\mu\text{m}$ | $3 \pm 2\%$ | Medium | $4 \pm 2\%$ | Low |

Together this setup forms a synthetic consortium where spatial interactions can be manipulated in a three-dimensional environment, and which allows the detection of growth and interactions using flow cytometry.

### Under glucose competition receivers cannot interact with producers ∼15 µm away

In the example above glucose could not diffuse from beads and each agarose bead acted as an individual compartment. In contrast, when glucose can diffuse from agarose beads the model predicted that the glucose concentration flattens close to the producer. In that case receivers at a distance of 15 µm from a producer in the same bead are exposed to similar glucose concentrations as receivers in beads without a producer (Figure 2C). If this prediction was correct, we expected that most receivers can grow when the global glucose concentration builds up, while in case of glucose competition the glucose concentration stays low and even receivers 15 µm away from a producer in the same bead should not be able to grow. We therefore incubated agarose beads in CDM, which allows glucose diffusion from beads.

Without competing glucose-consumers in the CDM outside the beads 75±7% of the beads showed growth and these beads had a high fluorescence/scatter ratio (Figure 2D, Table 2). This indicates growth of both receivers with and receivers without a producer in their bead. When we took the same beads but added competing glucose-consumers outside the beads, only 15±3% of the beads contained growth. In the beads where growth was observed the fluorescence/scatter ratio was low, indicating that only producers grew (Figure 2D, Table 2). These results are consistent with the model predictions and show that under glucose competition receivers cannot interact with producers even if they are only ∼15 µm away.

Without competing glucose-consumers still 25±7% of the beads were gated as “no growth”, although the model predicted that all receivers could grow (Figure 2C and 2D). These beads could be false negatives caused by our conservative gating strategy, or by empty beads with single fluorescent cells attached to their outside. Conversely, for the beads gated as “growth”, we observed an increased fluorescence/scatter ratio compared to the single receiver controls (Figure 2D). It is known that fluorescence of individual cells increases with decreasing growth rate [32, 33], suggesting that in co-cultures the higher fluorescence/scatter ratio could be caused by glucose limited and therefore slower growth of the receivers in the beads.

Together this data shows that competition for glucose in a three-dimensional environment prevents interactions at ∼15 µm distance, because the presence of competing public good-consumers leads to steep concentration gradients.

### Aggregated producers and receivers interact even under glucose competition

In the presence of steep concentration gradients microbial interactions might be facilitated by bringing producers and receivers in close proximity. Consistently, the model predicted that cell aggregation would allow receivers to grow under glucose competition (Figure 2E). We developed a protocol to make producer-receiver aggregates. Defined aggregates were formed by adding positively charged producers to an excess of negatively charged receivers, ensuring that producers were directly surrounded by receivers. In this way we obtained a mixture of single receivers and aggregates of one producer and approximately eight receivers (Figure 1B). We aimed to add an aggregate to ∼21% of the beads, but we could only roughly estimate the aggregate concentration in the mixture based on the added amount of positively charged cells. However, underestimating this percentage would not affect the results, as we only analyze agarose beads with growth after incubation (supplementary information, section 3).

We incubated the formed agarose beads in CDM with competing glucose-consumers and after incubation we saw an increased scatter in 3±2% of the beads (Figure 2F, Table 2), indicating only growth in beads with both producers and receivers. The fluorescence/scatter ratio of beads with growth was increased compared to the producer mono-culture (Figure 2F), indicating growth of both producers and receivers. Beads with grown aggregates of non-producers and receivers showed a fluorescence/scatter ratio similar to the producer mono-culture (Figure 2F), indicating that the aggregation protocol or the data analysis procedure did not influence the readout.

Therefore, the results show that cell aggregation facilitates microbial interactions, even in a three-dimensional system with competition for the public good.

### Aggregation results in dense micro-colonies, facilitating growth of receivers with low affinity and low V_max_ glucose transporters

To study the effect of glucose uptake efficiency on the receiver response, we modelled producer-receiver aggregates with receivers that have glucose-transporters with different affinities (K_m_) and maximal uptake rates (V_max_). Specifically, we modelled receivers that had one of the three different glucose transporters of *L. lactis* [22] (supplementary information, section 4.5). Within aggregates the effective diffusion coefficient (D_eff,s_) is described to be 10-70% of the diffusion coefficient in water (D_s_), depending on the aggregates’ density [29, 31]. When we set D_eff,s_ to10% of D_s_, the model predicts that receivers with the low K_m_ and high V_max_ transporter PTSman (K_m_ = 0.013 mM, V_max_ = 0.22 µmol/min/mg protein) consume about 90 times more glucose than receivers with a high K_m_ or low V_max_ transporter (PTScel: K_m_ = 8.7 mM, V_max_ = 0.25 µmol/min/mg protein, glcU: K_m_ = 2.4 mM, V_max_ = 0.08 µmol/min/mg protein) (supplementary information, section 4.5). When D_eff,s_ is 70% of D_s_, this difference is almost 350 fold.

As the model predicted less glucose consumption by receivers with low glucose affinities and low maximal glucose uptake rates, we constitutively expressed GFP in three engineered *L. lactis* NZ9000 strains which each contain only one of the three glucose transporters [27]. We subsequently analyzed if their uptake was high enough to interact with producers. As we saw before, the fluorescence/scatter ratio in mono-culture controls decreased with an increasing growth rate (Figure 3 and supplementary information section 5). The data further show that in producer-receiver aggregates even the low affinity receivers could grow and the differences in fluorescence/scatter ratio between the mutants were small (2-3 fold). Based on the model this data suggests that dense micro-colonies with a low D_eff,s_ were formed. Aggregates with receivers containing the low K_m_ and high V_max_ transporter PTSman showed the highest fluorescence/scatter ratio. Consistent with the model predictions this result suggests that PTSman containing receivers have the highest glucose uptake rate.

**Figure 3.**
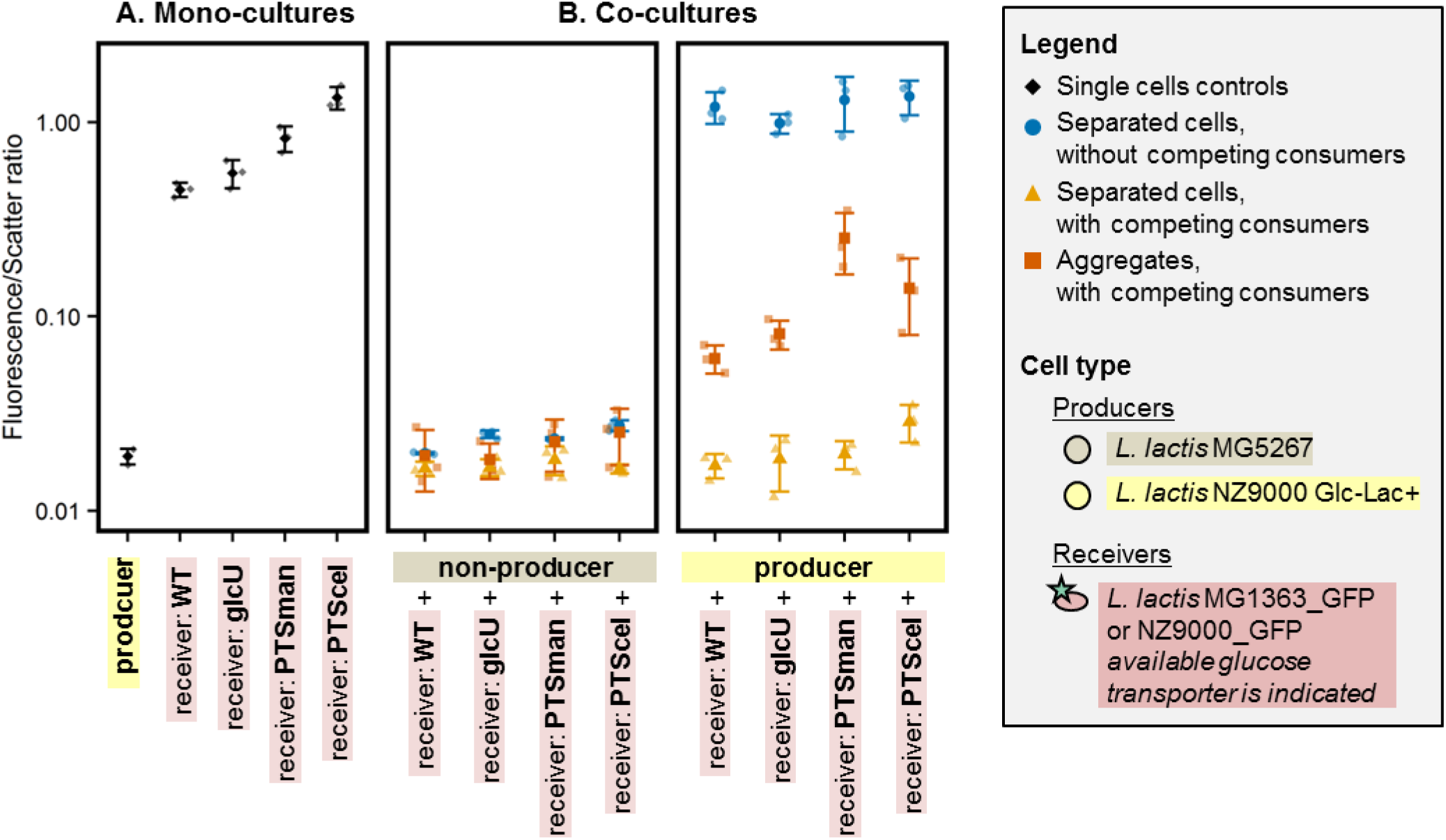
Effect of glucose transporter affinity and V_max_ on the receiver response. Mono- and co-cultures of (non-)producers and receivers with different glucose transporters were incubated in agarose beads surrounded by CDM. **(A)** Producer mono-cultures were incubated in presence of lactose, receiver mono-cultures in presence of glucose. Agarose beads contained separated cells. **(B)** Co-cultures were incubated in presence of lactose. Agarose beads contained either separated cells (∼15 µm between cells within a bead), or producer-receiver aggregates (0 µm between cells). The beads were incubated in CDM with and without competing glucose-consumers. For each culture the median and standard error of the median of the fluorescence/scatter ratio is shown (n=3). Receivers are ordered based on their growth rate (supplementary information section 5).

Altogether the data show that in a three-dimensional system with a metabolite consuming sink a steep concentration gradient is obtained, and cells only ∼15 µm away from each other cannot interact through glucose cross-feeding. This physical constraint can be overcome by bringing cells together in the low micrometer range, as achieved through cell aggregation - physical contact.

## Discussion

Contact-independent interactions can be local or global, depending on the profile of the concentration gradient. Previous studies quantified interaction distances either in monolayers of cells (two-dimensional system) or in absence of a competing metabolite-sink [13, 14]. While these studies give valuable insight, they have a limited resemblance to natural ecosystems, which are typically three-dimensional and harbor competing organisms and other metabolite-removing sinks. A reaction-diffusion model predicted that in three-dimensional systems the concentration gradient drops much faster than in two-dimensional systems (the maximal concentration is halved at 0.6 µm instead of 24 µm), suggesting that in natural ecosystems interaction ranges might be much shorter than the *in vitro* two-dimensional systems predict. To better understand how concentration gradient constraints affect the composition and fitness of natural communities, we analyzed the impact of cell-to-cell distance on unidirectional cross-feeding in three dimensions in the presence and absence of a competing metabolite consumer as a public good-removing sink.

Without competing glucose-consumers we observed a global response: receivers grew at all distances from producers (Figure 2D, Table 2). When we added competing glucose-consumers only receivers aggregated with producers could grow (Figure 2D and 2F, Table 2). Diener *et al.* observed a similar pattern of local and global interactions during *S. cerevisiae* mating [14]. Haploid cells secrete a peptide, which is sensed and degraded by haploid cells of the opposite mating type. This results in a local high peptide concentration and local interactions: cells from opposite mating types initiate mating specifically in each other’s direction. However, incubation of mutants that could not degrade the peptide resulted in a global high peptide concentration, and independent of their location cells initiated mating in different directions. For wildtype cells Diener *et al.* predicted that the maximum information content of the peptide distribution is similar for cells ∼17 and ∼2 µm away from each other [14], suggesting that yeast cells interact efficiently when they are 17 µm away from each other. Similar interaction distances (3.2-12.1 µm) were found by Dal Co *et al.* when they grew bidirectionally cross-feeding *E. coli* cells in a microfluidic chamber [13]. Our data however show that in a three-dimensional environment with a metabolite-sink the interaction distances are shorter, as producers and receivers ∼15 µm away from each other could not cross-feed (Figure 2D). Within aggregates cross-feeding was possible, but it was still less efficient than what was achieved in presence of high concentrations of the secreted metabolite (Figure 2B and 2F).

These results match the model prediction that interaction distances in three-dimensional systems are shorter than in two-dimensional systems. It furthermore indicates that the presence of a metabolite-sink also affects the interaction distance. In the set-ups of Dalco *et al.* and Diener *et al.* the metabolite is degraded/consumed only by the receiver itself, so the metabolite concentration will only decrease close to receiver cells. When receivers compete with other metabolite-sinks, such as competing metabolite consumers or a dilute system, the overall metabolite concentration will be lowered, resulting in shorter interaction distances. Indeed, Koschwanez *et al.* showed that at low cell densities and low sucrose concentrations, where the volume acts as the main metabolite-sink, *S. cerevisiae* cannot grow, even though invertase splits sucrose into glucose and fructose in the periplasmic space, so very close to the receiver cell [8].

In the presence of a competing metabolite-sink the concentration of the exchanged metabolite is low, and we therefore expected that variation in the receivers’ import affinity would influence the interaction efficiency. However, within aggregates we observed only small differences in growth of high and low affinity receivers (Figure 3), suggesting that dense micro-colonies with a low diffusion rate were formed (supplementary information, section 4.1). Aggregating cells therefore seem to kill two birds with one stone: they decrease both the cell-to-cell distance and the diffusion rate, two factors which were previously reported to promote interaction [15, 34]. The resulting increase of the local concentration might also explain why Koschwanez *et al.* saw that yeast cells which could not grow on low sucrose concentrations due to their low cell density, could grow when the same amount of cells was aggregated [8].

In our set-up the local glucose concentration increases with time, because the producers grew independently from the receivers. Future research could focus on bidirectional cross-feeding systems, where producer growth is limited by receiver growth. In that case the initial production rate will be lower and receiver affinity might play a bigger role.

Controlled metabolite exchange is a critical feature of living cells [35], and forms the basis for extracellular metabolism of nutrients and interactions with other cells. Concentration gradients constrain the distance over which these interactions can occur and it therefore shaped the evolution of the molecular mechanisms involved in these interactions. Slow diffusion of large, aggregated resources like particulate iron (>0.4 µm) can for instance cause cellular iron uptake to become diffusion limited. It is therefore hypothesized that cells secrete siderophores to form fast diffusing iron-siderophore complexes after iron dissolution, in this way increasing their iron uptake rate [36]. It is furthermore known that many extracellular substrate-degrading enzymes are attached to the cell, which places the source (enzyme) close to the receiver (cell). Invertase is for instance located in the periplasmic space of *S. cerevisiae* [37], the protease of *L. lactis* is attached to the cell wall [9] and in both fungi and bacteria cellulosomes are also attached to the cell wall [38]. Hauert *et al.* argue that when a producer also benefits from its own product, which is the case for extracellular enzymes, spatially structured localization of cells is only advantageous when the enzyme production costs are high [39]. Attachment of extracellular enzymes to the cell wall therefore suggests that these enzymes are costly, and indeed, Bachmann *et al.* showed in *L. lactis* that protease negative strains outcompeted protease positive strains with a cell wall bound protease, unless they were >1 mm apart (cell density of <10^3^ cells/mL) [9].

In the presence of a competing public good-sink interacting cells can aggregate, for example in biofilms, to reduce the diffusion distance and diffusion rate [40], which increases the efficiency of their interactions [15, 34]. During evolution of cooperation in which costly compounds are secreted, wildtype non-cooperators typically form such a competing public good-sink, indicating that cell-to-cell distances well below 15 µm are required to evolve costly cooperation. However, aggregation is not always increasing interaction efficiency, because it also slows down the diffusion of inhibiting metabolic end-products from the micro-colony and the diffusion of extracellular nutrients into the micro-colony. Aggregation of the cross-feeding yoghurt consortium (*Lactobacillus bulgaricus* and *Streptococcus thermophilus*) in 100-300 µm capsules reduced for instance their growth and acidification rates, and proteolysis was only faster in the first hour [41], indicating that in this case the aggregation costs did not outweigh the benefits. Aggregation also allows (evolution of) contact-dependent transfer mechanisms, like nanotubes or vesicle chains. To our knowledge *L. lactis* does not exchange cytosolic material using these contact-dependent transfer mechanisms and the model indicates that just diffusion can explain our experimental results.

Consequences of concentration gradient constraints are not limited to bacteria and yeasts. Plants, fungi and other (organisms with) large cells use intracellular concentration gradients to regulate amongst others cell polarity, cell division and cell size [42–44]. Although the rate of diffusion in the cytoplasm is fast, cells can use spatially structured protein modification systems as source or sink, to create concentration gradients [45, 46]. It is therefore important to think about the constraints - and opportunities - that concentration gradients may impose on cellular interactions, how it shaped their evolution and their role in microbial consortia, and how researchers can use these principles to understand and steer these processes.

## Acknowledgements

We thank Sieze Douwenga and Daan de Groot for fruitful discussions.

R.J.v.T., I.v.S., E.Z., J.A.H., O.P.K, B.T. and H.B. were financed by the Netherlands Organisation for Scientific Research (NWO), as part of the research programme TTW with project number 13858.

A.L. and T.R. were financed by the Slovenian Research Agency (Grant no. J4-7640, J1-9194, N1-0100), international grant supported by Helmholtz-Zentrum Dresden-Rossendorf and the European Commission (Grant no. 826312) and the European Regional Development Fund (Grant No. UIA02-228).

## Author contributions

R.J.v.T., B.T. and H.B. conceived the study, designed experiments, interpreted the data and wrote the paper. R.J.v.T., I.v.S. and E.Z. carried out the experiments. T.R. and A.L. developed the aggregation protocol. C.P. and R.J.v.T. built the COMSOL Multiphysics model. J.A.H. and O.P.K. constructed the strains *L. lactis* NZ9000_PTSman_GFP, *L. lactis* NZ9000_PTScel_GFP and *L. lactis* NZ9000_glcU_GFP. All authors helped improving the manuscript.

## Supplementary information

### Section 1: Agarose bead size- and volume-distributions

We prepared agarose beads surrounded by oil and made pictures with a microscope (9 per emulsion, Figure S1A shows an example). Pictures were subsequently analyzed with ImageJ to identify the beads (Figure S1B), and to measure size- and volume distributions (Figure S1C and S1D). Small droplets were not always identified, but as they contain only little volume this only marginally affects the analysis. Beads on the edge of the picture were excluded from the analysis. Formed emulsions were polydisperse but distributions of replicates were reproducible, with mean volume ± SEM of 26±2 pL (diameter of 37 µm).

**Figure S1.**
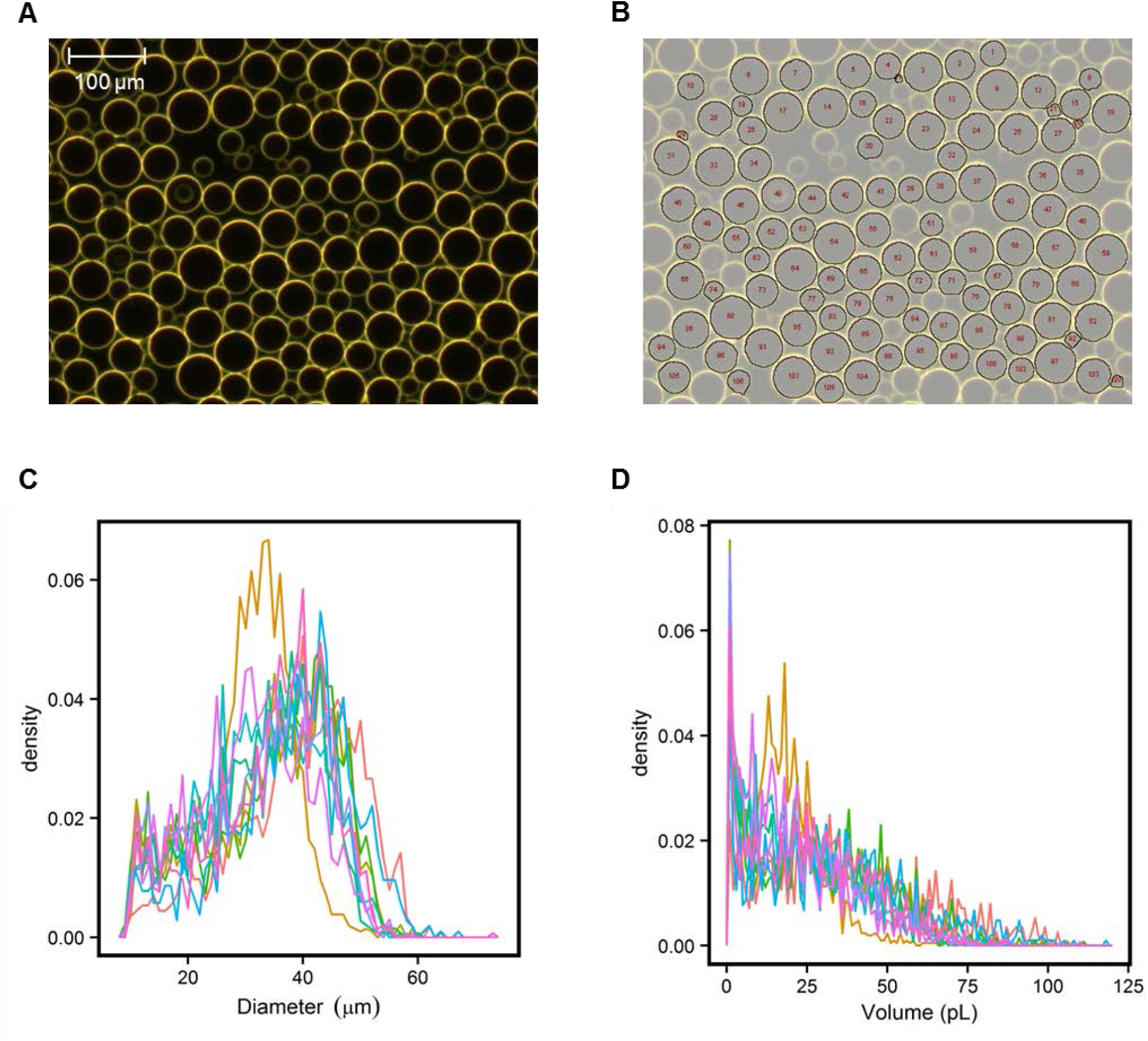
Agarose bead size and volume. **(A)** An example microscope picture of agarose beads. **(B)** Agarose beads identified in (A) after ImageJ analysis. **(C and D)** Agarose bead size (C) and volume (D) distribution (n=10 emulsions, 9 pictures per emulsion).

### Section 2: 10^9^ *L. lactis* MG1363 cells/mL as competing glucose-consumers

To establish if addition of 10^9^ *L. lactis* MG1363 cells per mL outside agarose beads prevented cross-talk between beads, we mixed beads with producers and beads with receivers and incubated them in presence of lactose in different spatial structures (Figure S2). After incubation surrounded by oil only producers were grown, which was expected as glucose could not diffuse from beads. When glucose could diffuse from beads and no *L. lactis* MG1363 cells were added outside the beads, both producers and receivers grew. However, in presence of 10^9^ *L. lactis* MG1363 cells per mL outside the agarose beads only producers grew, suggesting that glucose leaving beads with producers was mainly consumed by *L. lactis* MG1363 cells outside the beads and did not reach receivers in neighboring beads. The glucose concentration outside the beads probably did not exceed the low micro-molar range, as the K_m_ for glucose of the highest affinity transporter in *L. lactis* MG1363 is 13 µM [22] and the *L. lactis* MG1363 cell density was high.

**Figure S2:**
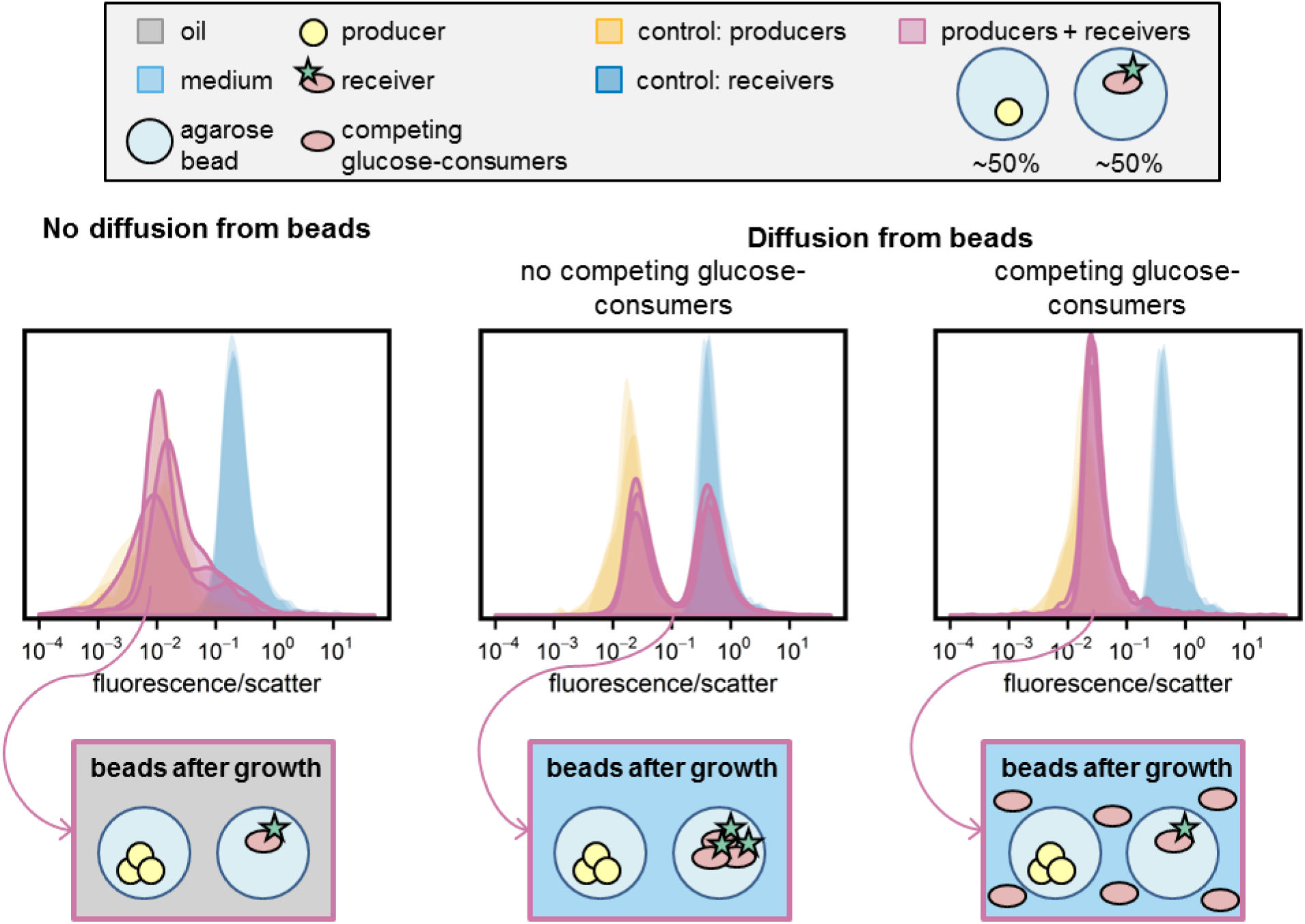
Addition of *L. lactis* MG1363 outside agarose beads prevents cross-talk between beads. Agarose beads with producers and agarose beads with receivers were mixed and incubated in three different spatial structures: surrounded by oil (no diffusion from beads, n=3), surrounded by CDM without *L. lactis* MG1363 (no competing glucose-consumers, n=3) and surrounded by CDM with 10^9^ single *L. lactis* MG1363 cells per mL (competing glucose-consumers, n=3). Histograms show the fluorescence/scatter ratio of the populations that were gated as “growth”.

### Section 3: Flow cytometry gating strategy and data analysis

Figure S3 shows the flow cytometry gating strategy and data analysis procedure.

**Figure S3.**
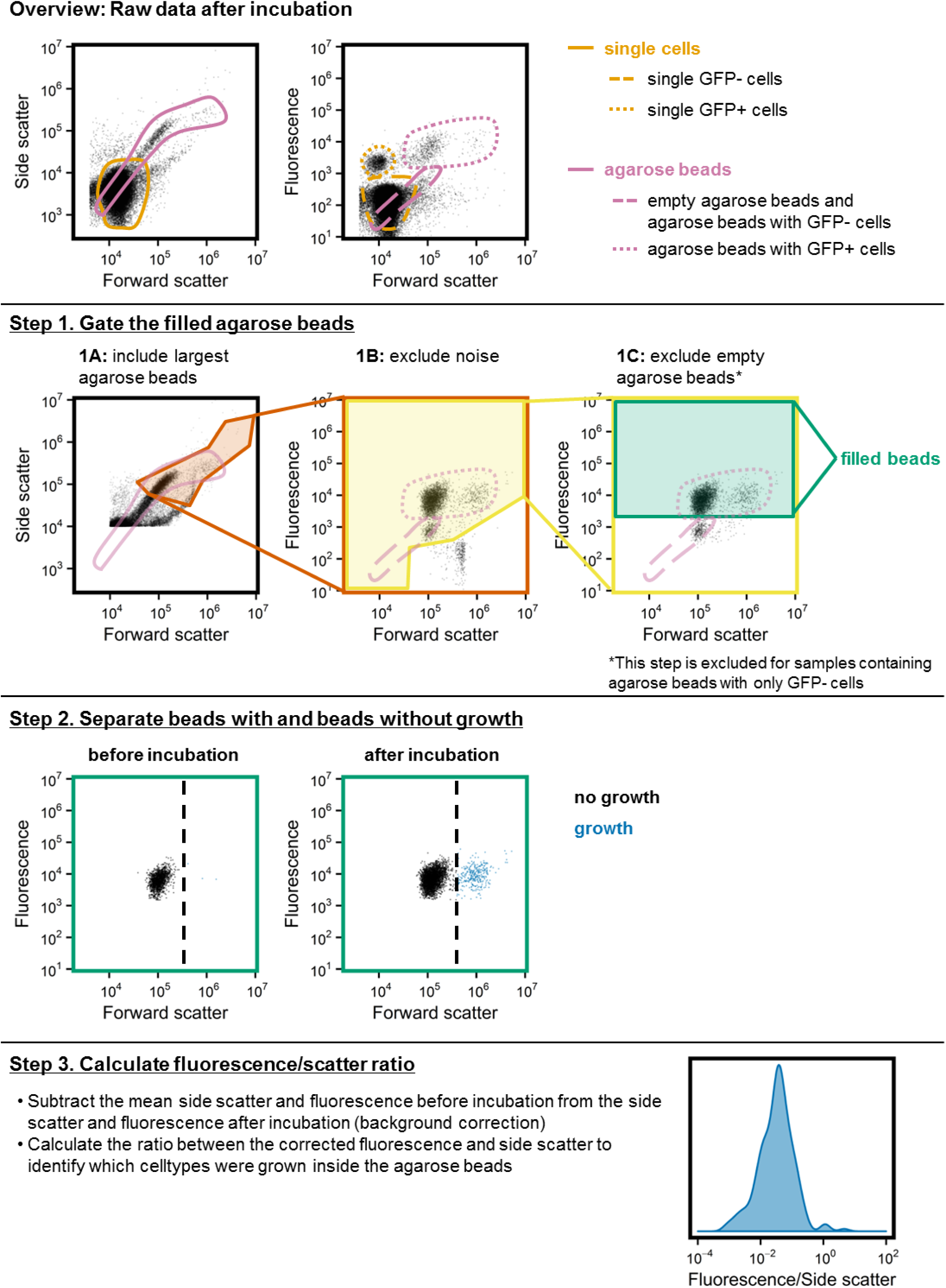
Flow cytometry gating strategy and data analysis.

#### Overview

Each agarose bead sample was measured twice. We first acquired an overview of the complete sample, containing both single cells and agarose beads. Drawn gates were based on measurements of empty agarose beads and single cells. Thereafter we used an increased forward and side scatter threshold to leave out most of the single cells, this data was used for gating and data analysis. **Step 1.** Beads containing cells (“filled beads”) were gated by including the largest agarose beads and excluding the noise and the empty agarose beads. For the control samples with only producers empty beads were not excluded, because before incubation agarose beads with GFP-producers could not be separated from empty beads. **Step 2.** Beads with and without growth were separated with a forward scatter threshold. This threshold was set for each sample individually, based on the forward scatter before incubation. This stringent gating might underestimate the amount of beads with growth, but it ensures that beads without growth are excluded from analysis. **Step 3.** The side scatter and fluorescence were background-corrected based on their values before incubation. The distribution of fluorescence/scatter ratios of background-corrected data is plotted to identify which cell-types were grown inside the agarose beads.

### Section 4: Reaction-diffusion model in COMSOL Multiphysics

#### 4.1 Geometry of the agarose bead model

Figure S4 shows the geometry of agarose bead model.

**Figure S4:**
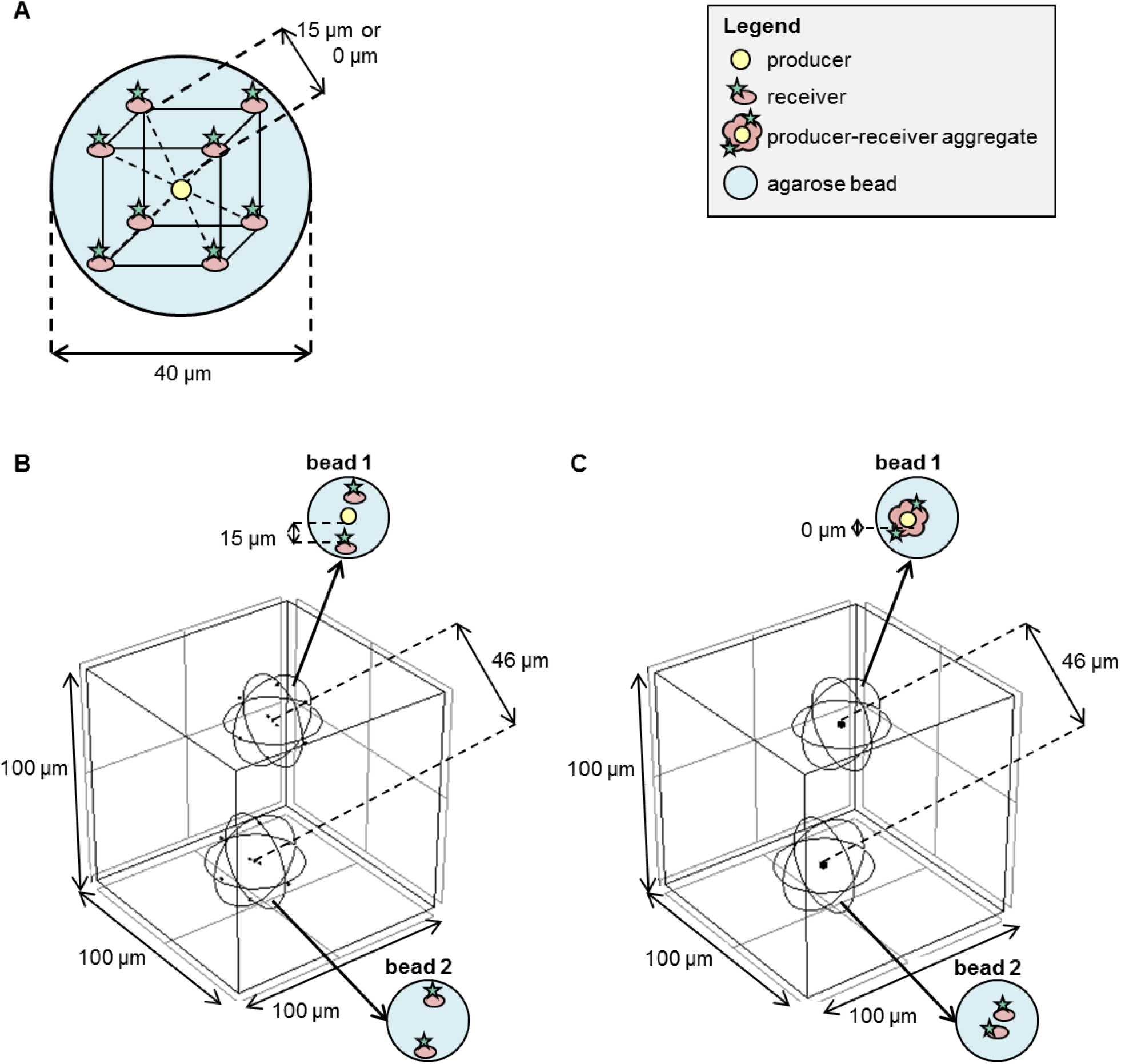
Model geometry implemented in COMSOL Multiphysics. **(A)** An agarose bead (sphere with diameter of 40 µm) contains 8 receiver cells and 0 or 1 producer cells. Each cell is a sphere with a diameter of 1 µm. Within the agarose bead receivers are placed at the virtual corners of a cube, and the producer in the middle. The distance between the surface of producers and receivers is either 0 or 15 µm. **(B and C)** An agarose bead with one producer (bead 1) and an agarose bead without a producer (bead 2) are placed in a cube of 100 µm (corresponding to 2·10^6^beads/mL). In (B) the distance between the producer and receiver is 15 µm, in (C) they are in contact (0 µm). Producers and receivers in contact are placed in a micro-colony (sphere with a diameter of 4 µm) with a reduced diffusion coefficient (D_eff,s_).

#### 4.2 Reaction-diffusion model and parameter values

##### Material balance

The spatial distribution (x,y,z) and change in time (t) of the concentration C_s_ (mol/m^3^) of glucose in the bead and surrounding liquid resulted from solving the partial differential equation which balances the diffusion rate with a reaction rate r_s_:

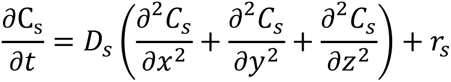

##### Diffusion

The same diffusion coefficient of glucose, D_s_ (m^2^/s), was used outside and inside the agarose beads. It was set to the value of D_s_ in water [29], because D_s_ in agarose gels is similar to that in water [30]. The effective diffusion coefficient in micro-colonies depends on the void fraction, i.e. volume not occupied by cells per total micro-colony volume, and the tortuosity [13, 40]. Cells growing in an agarose matrix form dense colonies, therefore the effective diffusion coefficient within the colonies (D_eff,s_) was set to 10% of the diffusion coefficient in water (D_s_) [29, 31].

##### Reaction

The net glucose rate r_s_ (mol/m^3^/s) results as the difference between production and consumption at a certain position in space, r_s_ = q_p_C_x_ - q_s_C_x_. The specific glucose production rate (q_p_) of *L. lactis* NZ9000 Glc-Lac+ is the same as its specific lactose uptake rate, as each lactose molecule contains one glucose molecule. The q_p_ was therefore set to a constant value of 1 molP/CmolX/h [47] and applied within the producer cells. Simulations which did include the lactose concentration and Monod kinetics for lactose consumption yielded similar results as simulations with a constant q_p_, therefore we adopted the simpler constant rate. For receivers the glucose uptake was assumed with a saturation (Monod) kinetics, q_s_ = q_s_^max^·C_s_/(K_s_ + C_s_). We used the K_s_ of the highest affinity glucose transporter of *L. lactis* MG1363 [22], and q_s_^max^ was set to 1 molS/CmolX/h [48]. To calculate the biomass concentration C_x_ (CmolX/m^3^) we assumed a molecular weight of biomass of 24.6 grams per Cmol dry biomass (CH_1.8_O_0.5_N_0.2_) [49, 50], a cellular water content of 70 wt% [51] and a cellular density of 1000 g/L [51]. These values lead to a glucose production or maximal glucose consumption rate of 38 mol/m^3^/s. A competing glucose-consumer was modelled by adding glucose consumption outside agarose beads with the same Monod kinetics as that of receivers. Cell growth was not incorporated in the model.

##### Boundaries

The concentration at the agarose bead surface was based on a partition coefficient which was set to 0 when incubation in oil was modelled, and to 1 for incubation in CDM. The liquid domain (cube) boundaries were insulated (no-flux boundary condition).

##### Parameters

Table S1 lists the default parameters used in the COMSOL Multiphysics model with sources for their values.

**Table S1:**
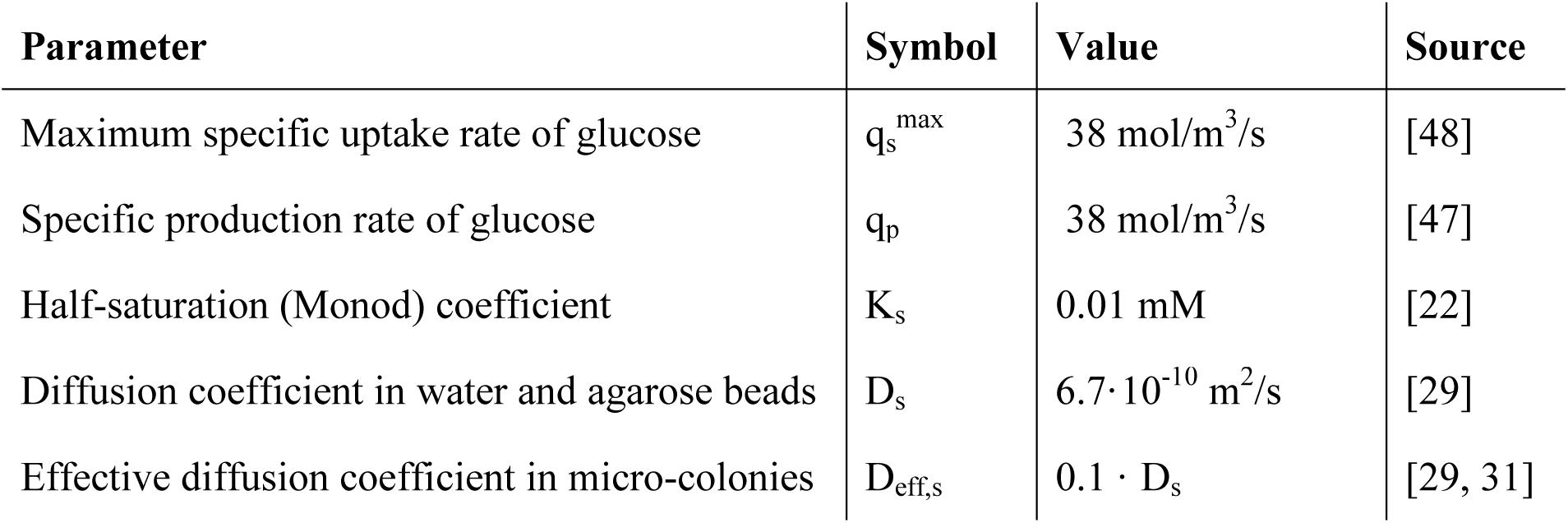
Default parameter values of the COMSOL Multiphysics model

#### 4.3 Predicted concentration gradients in two- and three-dimensional reaction-diffusion systems

To analyze the difference in concentration gradients in two- and three-dimensional systems we modelled production by one producer cell in two different geometries, as represented in Figure S5. In the two-dimensional system the model predicts that the product concentration is halved at 24 µm from the producer, whereas in the three-dimensional system this happens at only 0.6 µm.

**Figure S5:**
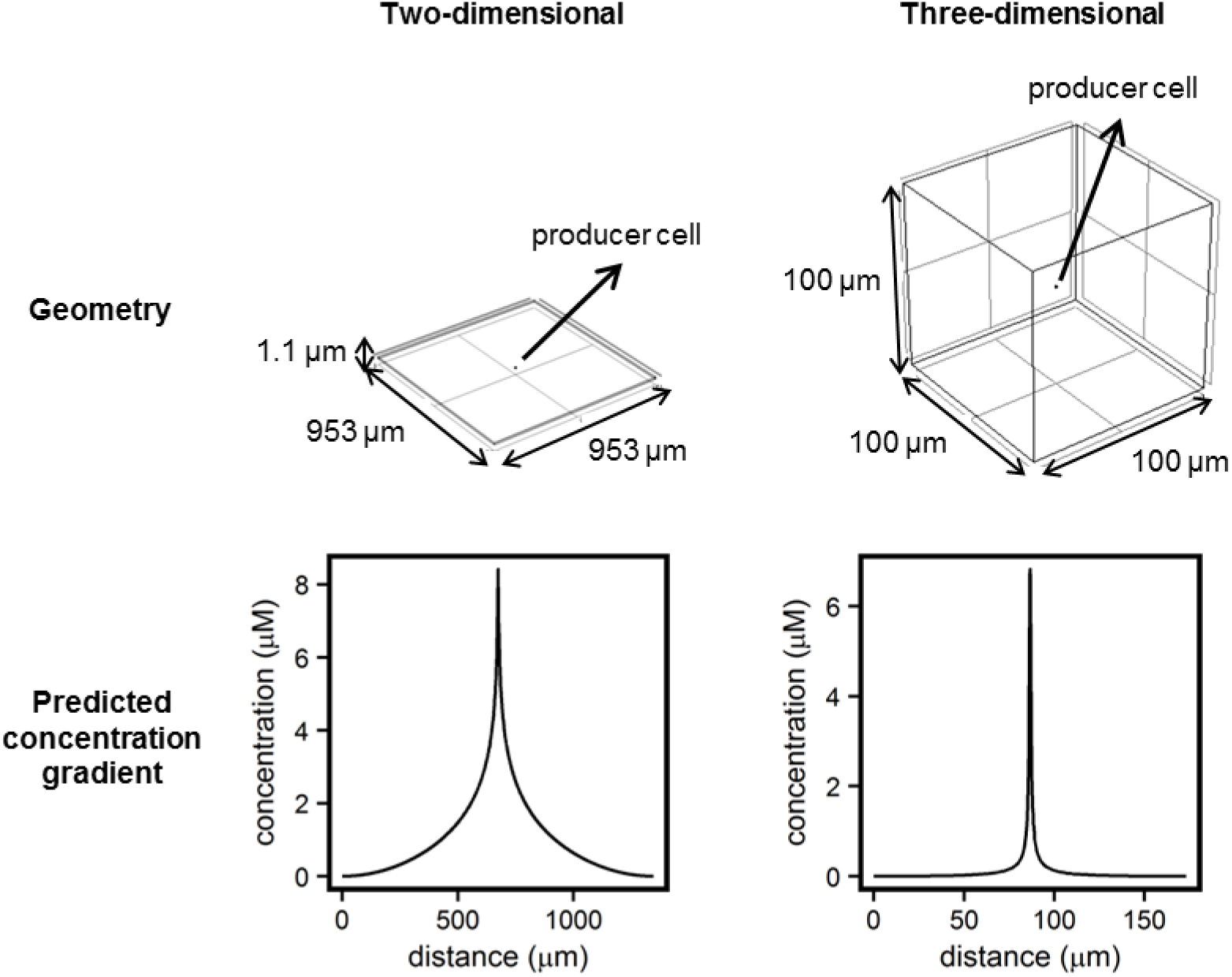
Predicted concentration gradients in two- and three-dimensional reaction-diffusion systems. **A** producer cell with a diameter of 1 µm was placed in the middle of a very thin rectangular block (1.1 µm thickness) to represent a quasi-two-dimensional system. For the three-dimensional system it was placed in the middle of a cube. For the cube, concentrations at the cube boundaries were set to zero. For the thin plate the concentrations at the four lateral faces were set to zero, and the top and bottom boundaries were insulated (no-flux boundary condition). The total volume of both systems was 1 nL (1·10^6^ cells/mL). A time dependent study in COMSOL Multiphysics yielded concentration gradients at several moments. The figure shows the concentrations along the diagonal after 5 hours.

#### 4.4 Sensitivity analysis for q_p_ and q_s_^max^

Figure S6 shows the predicted glucose concentration (Figure S6A) and glucose production rate (Figure S6B) over a plane crossing the producer cell and four of the eight receiver cells. Profiles for aggregated cells and for cells ∼15 µm away from each other are shown. Because it is difficult to know the actual q_p_ and q_s_^max^ inside agarose beads, we also performed a sensitivity analysis (Figure S7). A 5-fold change in q_p_ and q_s_^max^ resulted in similar concentration gradients and it did not affect our hypotheses.

**Figure S6:**
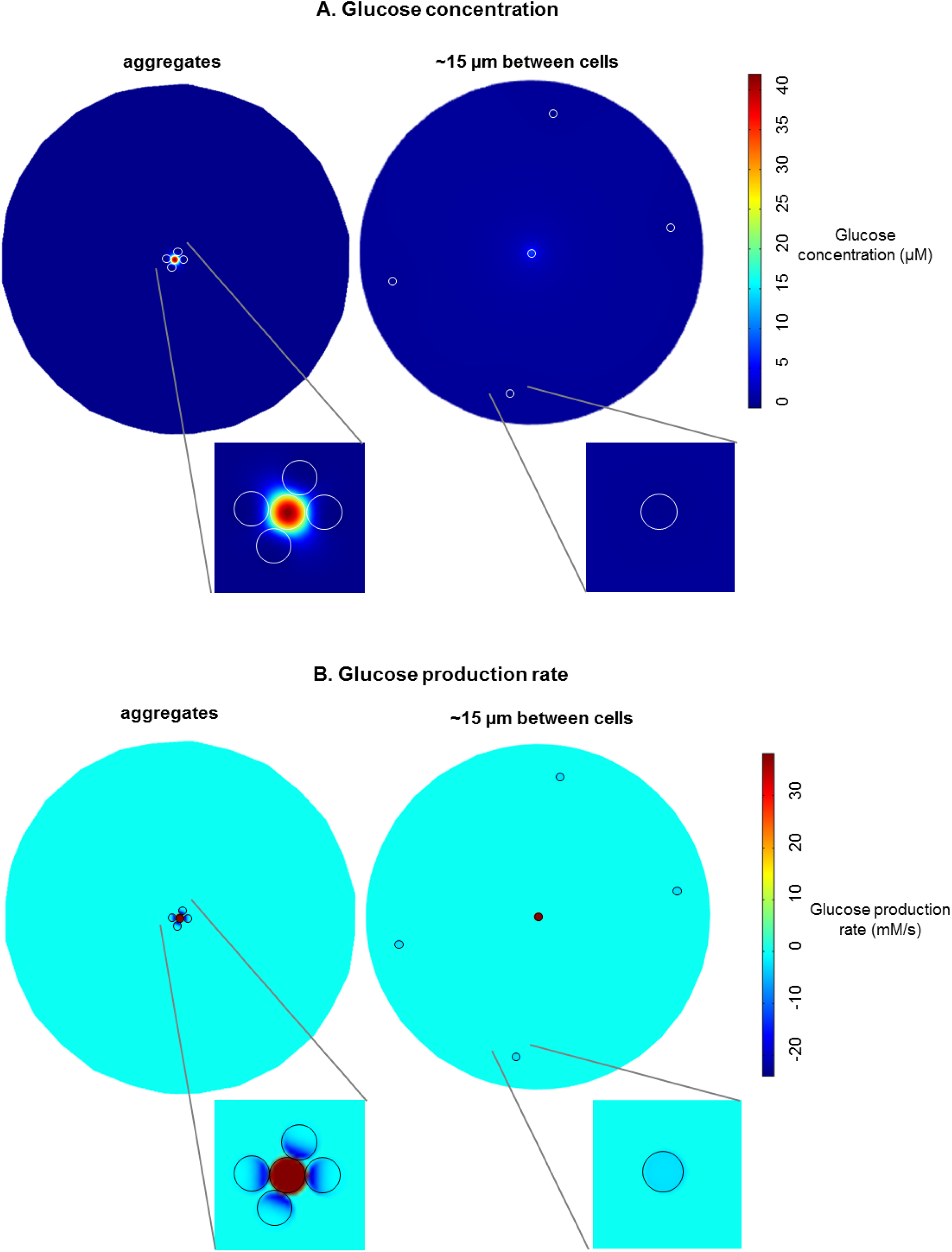
Predicted glucose concentration (A) and glucose production rates (B) in agarose beads surrounded by CDM. The glucose concentration **(A)** and the glucose production rate **(B)** are plotted over a plane crossing the producer cell and four of the eight receiver cells. Profiles for aggregated cells and for cells ∼15 µm away from each other are shown.

**Figure S7:**
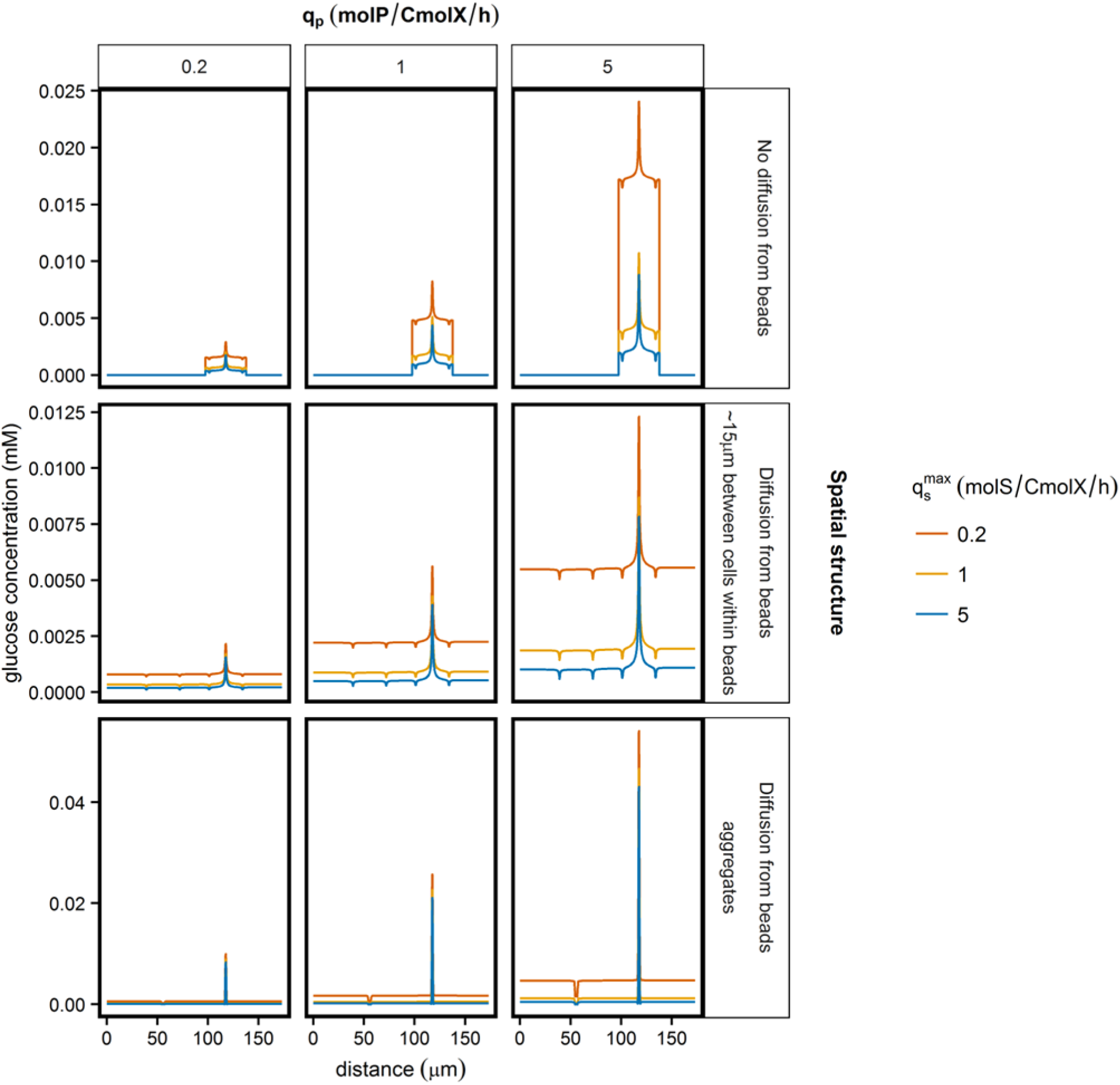
Predicted concentration gradients after varying q_p_ and q_s_^max^. q_p_ and q_s_^max^ were increased and decreased five-fold and the effects on the predicted concentration gradients in different spatial structures are shown.

#### 4.5 Model predictions for the glucose uptake of receivers with different affinities

We analyzed the effect of the glucose affinity of receivers on their ability to utilize the glucose made by the producer. To model the individual glucose transporters of *L. lactis* MG1363 in COMSOL the K_s_ values as reported by Castro *et al.* were used [22]. For *L. lactis* NZ9000_GFP_glcU the q_s_^max^ was reduced with a factor four, which reflects the differences in V_max_ of the transporters [22]. We calculated the glucose uptake for the different mutants after 5 hours in presence of competing glucose-consumers, without considering growth of the cells (Figure S8). The effective diffusion coefficient (D_eff,s_) varies from 10-70%, depending on the density of the micro-colony [29, 31]. Figure S7 shows the glucose uptake when D_eff,s_ is 10%, 30% and 70%. We included a sensitivity analysis for five-fold changes in q_p_ and q_s_^max^ values, which all showed similar trends as the reference (1x q_p_ and 1x q_s_^max^).

**Figure S8:**
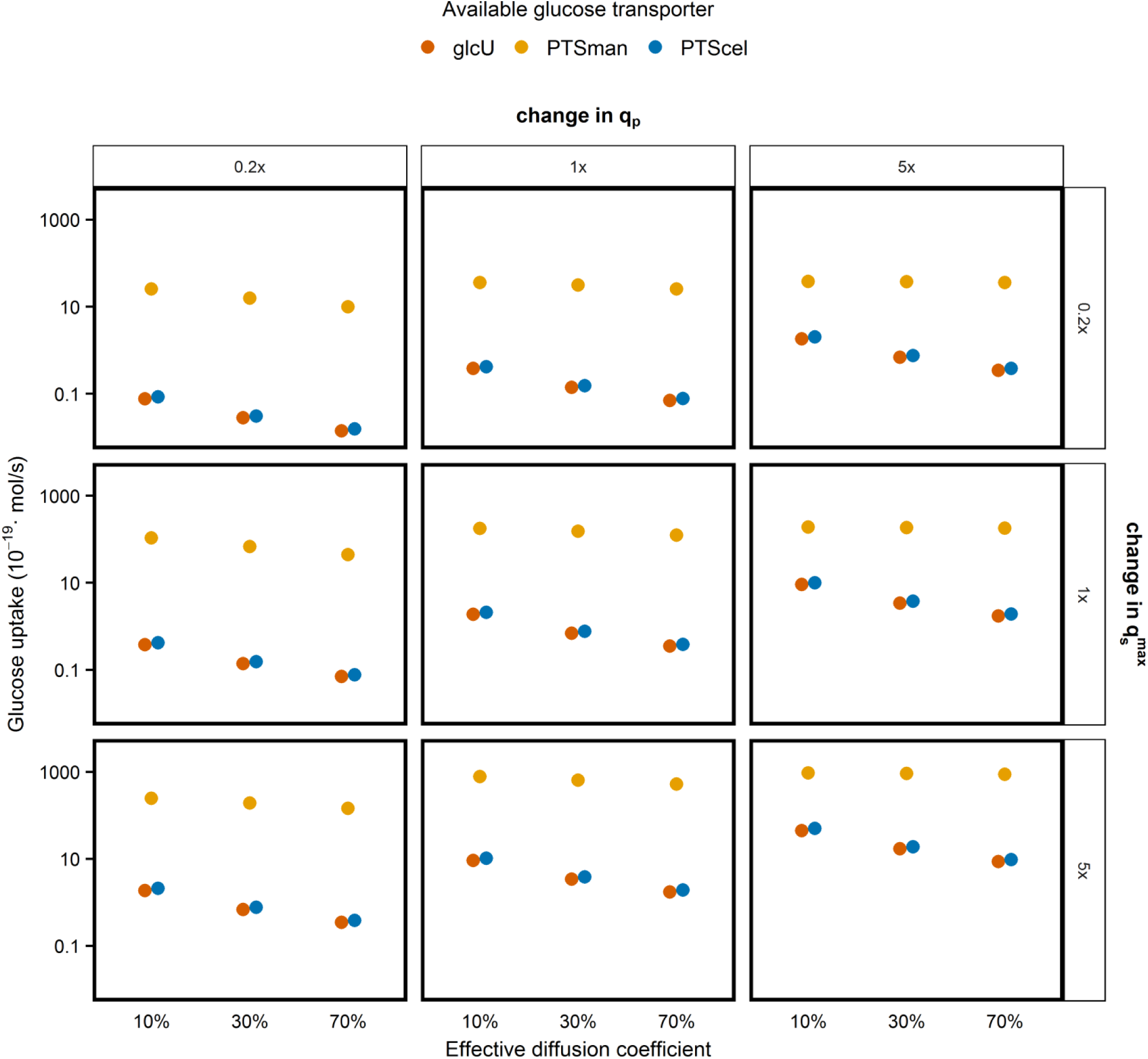
Predicted glucose uptake. We modelled receiver cells with different glucose affinities and calculated the predicted glucose uptake. A sensitivity analysis for five-fold changes in q_p_ and q_s_^max^ values was incorporated. Each simulation contained eight receiver cells aggregated with a producer (Figure S5), this figure shows the combined glucose uptake of these receivers.

### Section 5: Growth rate determination

Strains were incubated in CDM + 0.2 wt% glucose in a 96-well plate. The OD_600_ was measured every six minutes for 24 hours using a SPECTRAmax 384 plus plate reader (Molecular Devices, San Jose, CA, USA). OD_600_ measurements were background corrected, ln-transformed and the slope of the region with exponential growth was calculated as the growth rate (Table S2).

**Table S2.** Growth rates of receivers with different glucose transporters (n=22).

| Strain | Growth rate ± SD<br>(h<sup>-1</sup>) |
| --- | --- |
| *L. lactis* MG1363_GFP | 1.25 ± 0.04 |
| *L. lactis* NZ9000_GFP_glcU | 0.79 ± 0.04 |
| *L. lactis* NZ9000_GFP_PTSman | 0.74 ± 0.11 |
| *L. lactis* NZ9000_GFP_PTScel | 0.57 ± 0.03 |

### Section 6: Flow cytometry data for receivers glucose transporters with different affinities and V_max_

We constitutively expressed GFP in three previously constructed *L. lactis* NZ9000 mutants with a single glucose transporter [27], and analyzed their growth in different spatial structures. This experiment focused on beads incubated in CDM (allowing glucose diffusion from beads), as we expected that under these conditions the transporter characteristics of receivers would be important. Figure S9A shows the experimental results when receivers were ∼15 µm from a producer within the same bead and incubated in CDM, whereas in Figure S9B the beads were incubated in medium with 10^9^ glucose-consumers per mL. Figure S9C shows the experimental results of producer-receiver aggregates, incubated in CDM with 10^9^ glucose-consumers per mL. Without a competing glucose-consumers we observed growth of both receivers with and receivers without a producer in their bead (Figure S9A), while with competing glucose-consumers only producers could grow (Figure S9B). In producer-receiver aggregates receivers were able to grow, despite the presence of competing glucose-consumers (Figure S9C). The results were similar for all glucose transporters, and were consistent with the results of the wild-type (Figure 2, Table 2).

**Figure S9A.**
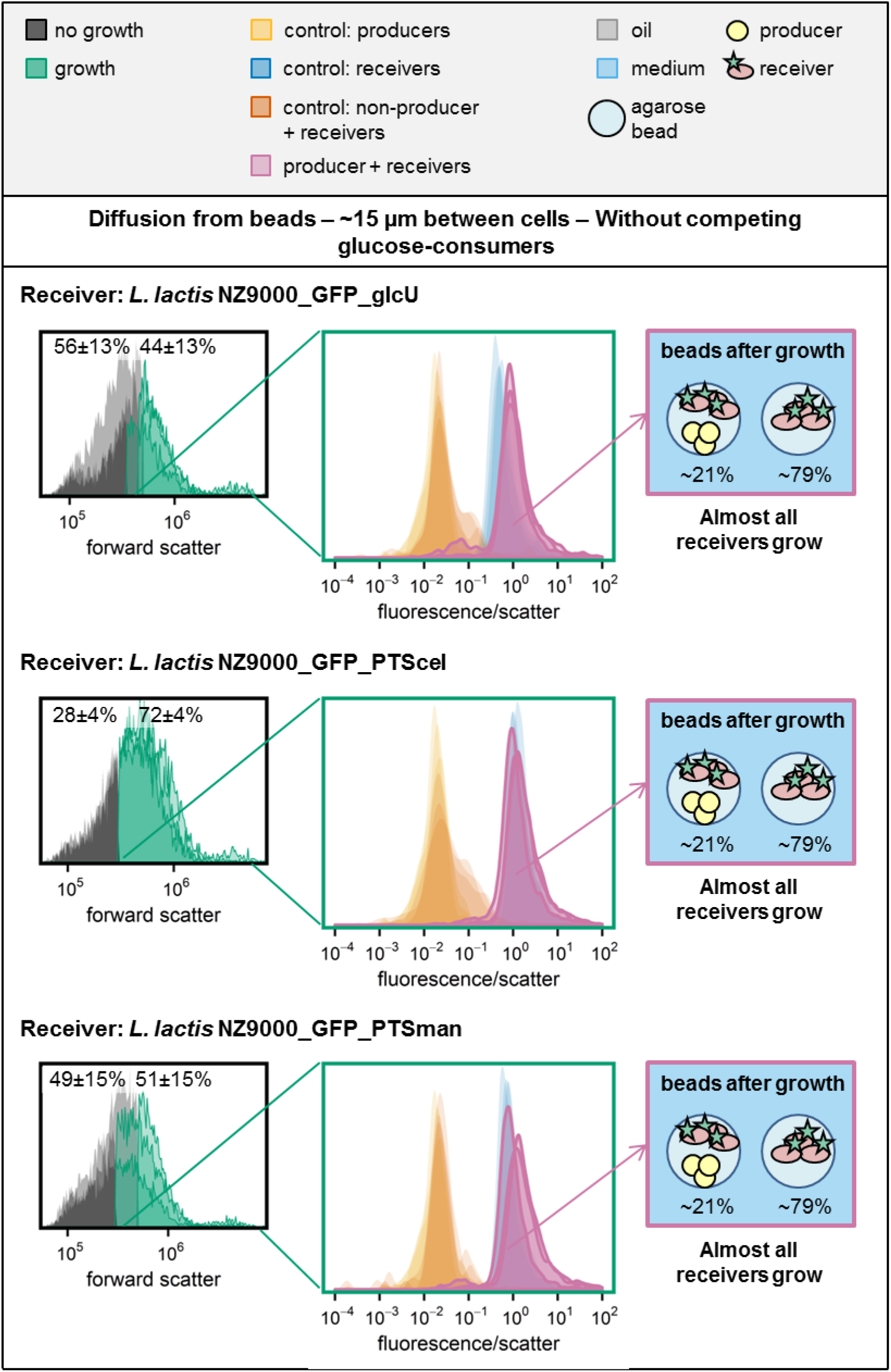
Glucose accumulates, allowing receivers to grow independent of the available glucose transporter. See complete caption on page 24.

**Figure S9B.**
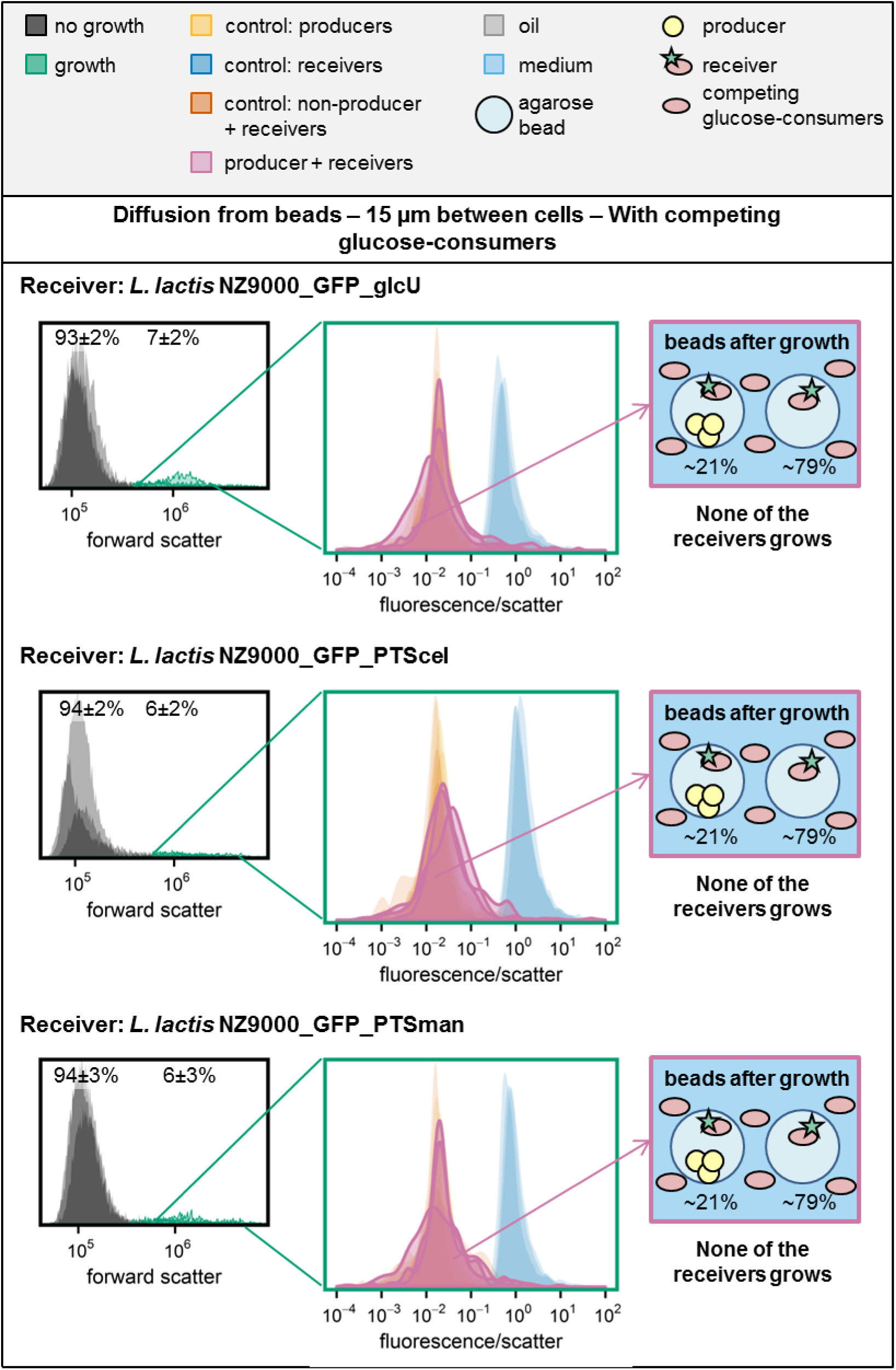
In presence of competing glucose-consumers receivers cannot grow, independent of the available glucose transporter. See complete caption on page 24.

**Figure S9C.**
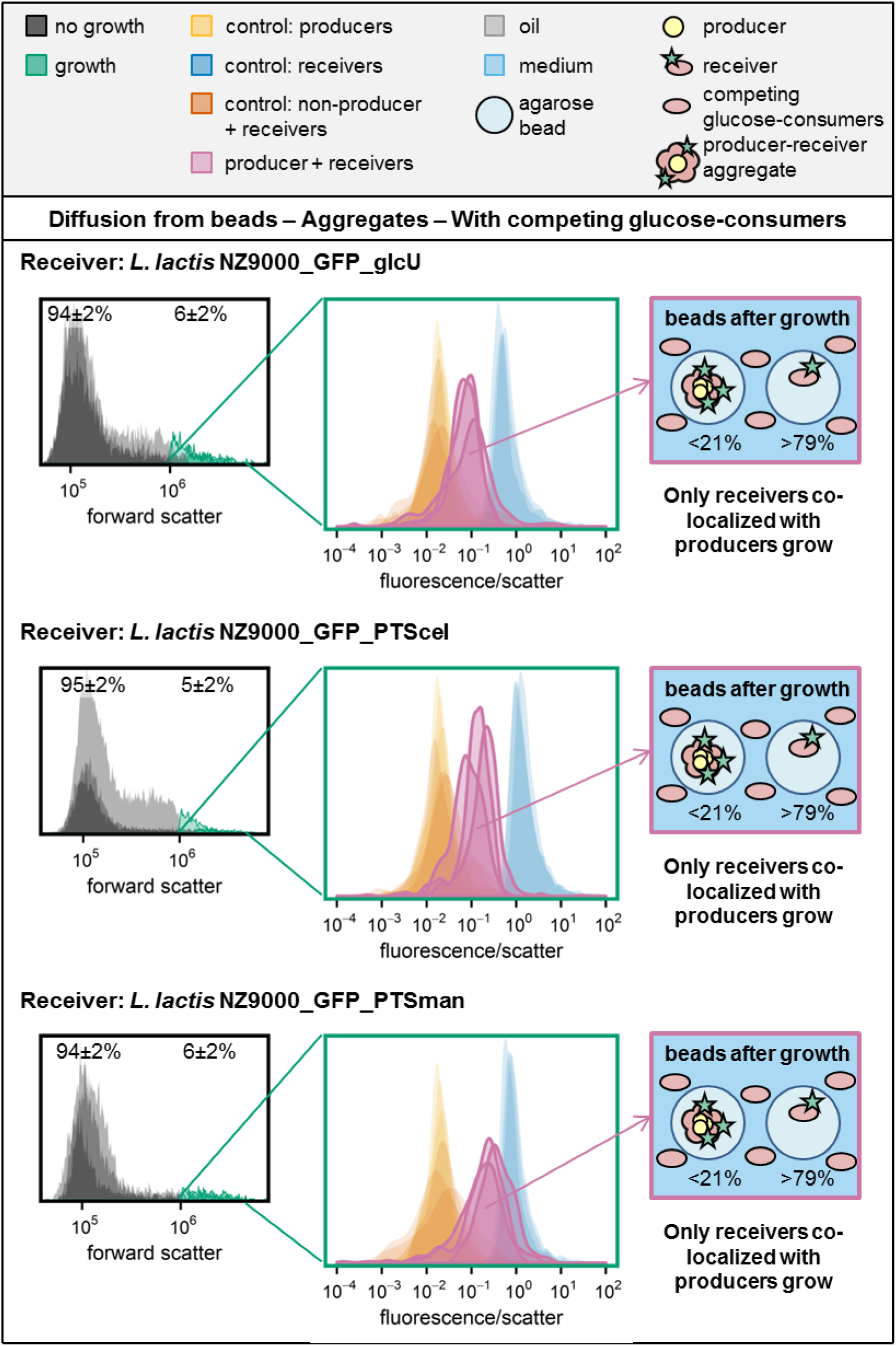
Despite the presence of competing glucose-consumers receivers within producer-receiver aggregates can grow, independently of the available glucose transporter. See complete caption on page 24.

**Figure S9: Response of consortia containing receivers with different glucose affinities.** The forward scatter histograms show the percentage of beads that were gated as “growth” in the co-culture of producer and receivers. The fluorescence/scatter histograms show the fluorescence/scatter ratio of beads that were gated as “growth” (n=3). Next to the co-culture of producer and receivers several control samples are included: receivers only, producers only and co-cultures of non-producers and receivers (n=3 for each of them). The non-producers and receivers, and the producers only controls are overlapping in all plots. The schematic drawing at the right shows the situation after growth, based on data from the two histograms. **Different panels contain different spatial structures: (A)** Receivers ∼15 µm from a producer within the same bead, incubated in CDM. **(B)** Receivers ∼15 µm from a producer within the same bead, incubated in CDM with 10^9^ glucose-consumers per mL. **(C)** Aggregates of producers and receivers, incubated in beads surrounded by CDM with 10^9^ glucose-consumers per mL.

